# *BundleCleaner*: Unsupervised Denoising and Subsampling of Diffusion MRI-Derived Tractography Data

**DOI:** 10.1101/2023.08.19.553990

**Authors:** Yixue Feng, Bramsh Q. Chandio, Julio E. Villalón-Reina, Sophia I. Thomopoulos, Himanshu Joshi, Gauthami Nair, Anand A. Joshi, Ganesan Venkatasubramanian, John P. John, Paul M. Thompson

## Abstract

We present *BundleCleaner*, an unsupervised multi-step framework that can filter, denoise and subsample bundles derived from diffusion MRI-based whole-brain tractography. Our approach considers both the global bundle structure and local streamline-wise features. We apply *BundleCleaner* to bundles generated from single-shell diffusion MRI data in an independent clinical sample of older adults from India using probabilistic tractography and the resulting ‘cleaned’ bundles can better align with the atlas bundles with reduced overreach. In a downstream tractometry analysis, we show that the cleaned bundles, represented with less than 20% of the original set of points, can robustly localize along-tract microstructural differences between 32 healthy controls and 34 participants with Alzheimer’s disease ranging in age from 55 to 84 years old. Our approach can help reduce memory burden and improving computational efficiency when working with tractography data, and shows promise for large-scale multi-site tractometry.

## 1 Introduction

Diffusion MRI (dMRI) allows us to examine white matter (WM) pathways in vivo and structural connectivity in the brain. Whole-brain tractograms (WBT), composed of streamlines, are used to model WM and can be generated using various tractography methods. However, inferring the geometry of neural pathways indirectly from water diffusion using tractography has limitations in practice: bundles in the brain often intersect, making it difficult to accurately infer long-range projections and determine termination points near the cortex [13]. In practice, WBTs may typically contain over 1 million streamlines per subject, including a high percentage of false positives (streamlines that do not represent true anatomy), and these also persist after bundle segmentation [27]. This can cause problems for downstream tasks, such as tractometry, where detected signals can be due to artifacts instead of true group differences. Many factors can affect the proportion of false positive streamlines – the quality of the incoming imaging data, the spatial and angular resolution of the acquisition, the reconstruction method for estimating the fiber orientation distribution function (fODF), and the tractography and bundle segmentation methods. This is a major concern when the dMRI data is collected in the single-shell low *b*-value regime. Large-scale multi-site neuroimaging consortia, such as ENIGMA, have processed large amounts of dMRI data most of it being single shell, low *b*-value data (*b*=600 − 1200*s/*mm^2^) [12], [16]. Post-hoc bundle filtering techniques become especially important in this context to reduce the proportion of spurious fibers that can affect the statistical analysis at the bundle level.

To address post-hoc filtering of tractography data, deep learning methods, such as an autoencoder [17] and geometric models [4], have been proposed for direct streamline filtering in WBT, where each streamline used in training is labeled as either valid or invalid. However, these models evaluate each streamline independently and are trained on reliably labeled WBTs instead of segmented bundles. Spurious fibers can occur in bundles where the streamline itself is structurally plausible but deviates from the overall fiber bundle shape. FiberNeat [6], filters streamlines in the low-dimensional embedding space using dimensionality reduction methods *t*-SNE and UMAP. The input to *t*-SNE and UMAP is a pairwise streamline distance matrix, which does not take into account the full structure of streamlines. In addition, both methods apply filtering by assigning ‘valid’ and ‘invalid’ labels to each streamline. However, when a streamline deviates only slightly or contains structural anomalies in a local neighborhood, it could potentially be addressed with smoothing instead of removal.

In this study, we propose *BundleCleaner*, an unsupervised multi-step framework to filter, denoise, and subsample bundles using both point cloud and streamline-based methods. This approach considers the global bundle structure as well as local streamline-wise features, and can be applied to bundles of varying shapes and proportions of erroneous streamlines. We applied *BundleCleaner* to bundles generated from single-shell dMRI in an independent cohort from India, and show that the cleaned bundles can improve alignment with the atlas bundles and reduce overreach. In a downstream tractometry analysis conducted using BUndle ANalytics (BUAN)[5], the cleaned bundles, represented with less than 20% of the original set of points, can robustly identify group differences in microstructural measures, with reduced memory burden and increased efficiency. Code for *BundleCleaner* is publicly available at https://github.com/wendyfyx/BundleCleaner.

## 2 Materials

We analyzed 3T single-shell dMRI data of the human brain in a pilot sample of 66 participants from the NIMHANS (National Institute of Mental Health and Neuro Sciences) cohort (mean age: 67.1 ± 7.4 years; 26F/40M). Thirty-four participants were diagnosed with Alzheimer’s disease (AD) and 32 were cognitively normal controls (CN). The dMRI data were acquired using a single-shell, diffusion weighted echo-planar imaging sequence (TR=7441 ms, TA = 630 s, TE=85 ms, voxel size: 2 × 2 × 2mm^3^, 64 slices, flip angle= 90^*◦*^, field of view (224mm)^2^). The protocol consisted of 64 diffusion-weighted (*b*=1000*s/*mm^2^), and 1 *b* = 0*s/*mm^2^ volume, where diffusion weighting was encoded along 64 independent orientations. Transverse sections of 2 mm thickness were acquired parallel to the anterior commissure-posterior commissure line. dMRI data were preprocessed to correct for artifacts with DIPY [9] and FSL [14], including noise [18], Gibbs ringing [19] [15], susceptibility induced distortions [1], eddy currents [2] and bias field inhomogeneity [26]. Whole brain tractograms were generated using constrained spherical convolution (CSD) [25] and probabilistic particle filtering tracking (PFT) [11] with the following parameters -8 seeds per voxel generated from the WM mask, step size of 0.2mm, angular threshold of 30^*◦*^, and the continuous map stopping criterion. RecoBundles[10] was used to extract 38 bundles for each subject in both native and MNI (Montreal Neurological Institute) space using a standard atlas in MNI ICBM 2009c space [28].

## 3 BundleCleaner

In developing filtering or denoising methods for white matter bundles, it would be beneficial to design methods that consider the sequential information within streamlines. However, some outliers can be determined by local features instead of all points on the streamline. It is important to design a smoothing algorithm that considers the global cohesion of the bundle as well as the smoothness of local streamlines. With this in mind, *BundleCleaner* contains the following steps:

– **Step 1**: Streamline resampling and pruning using QuickBundles [8]
– **Step 2**: Point cloud-based smoothing using Laplacian regularization [24];
– **Step 3**: Streamline-based smoothing using the Savitzky–Golay filter [23];
– **Step 4**: Streamline pruning and subsampling using QuickBundles (similar to step 1)

Methods used in each step are detailed below, and all parameters are listed with their default values in Table 1.

**Table 1.**
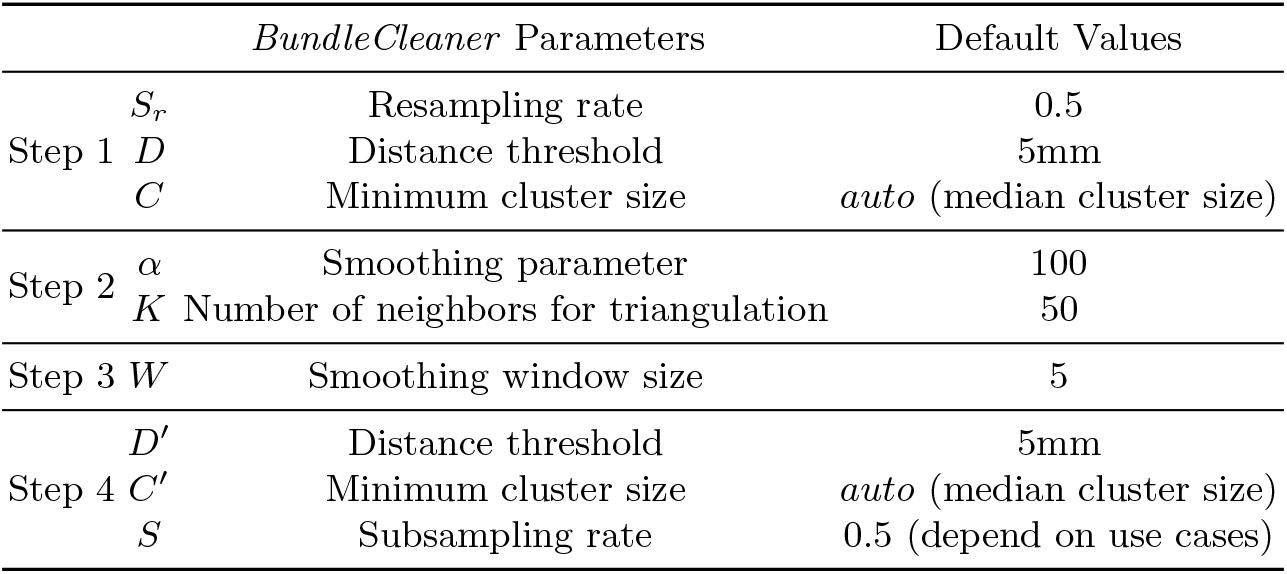
*BundleCleaner* parameters and their default values.

### 3.1 Step 1: Streamline Resampling and Pruning

Each bundle is initially loaded as a set of streamlines or 3D point sequences. Step size during fiber tracking is often smaller than the voxel size to produce more accurate streamlines [20], resulting in a dense representation. However, we can adequately represent streamlines with fewer points than those directly produced by the tractography [21]. To reduce the computational cost in Step 2 while maintaining the fixed step size, each streamline in the bundle is first resampled to *S*_*r*_% points equally spaced along the line.

As bundles with high levels of noise can be difficult to smooth directly, an initial step of pruning is applied using QuickBundles [8], an efficient streamline-based clustering method with a distance threshold *D*. The minimum direct flip distance (MDF) is used in clustering since streamlines are flip-invariant. Starting from one initial cluster, each streamline whose MDF distance to any existing centroid is smaller than *D* is added to the corresponding cluster; otherwise, it is added to a new cluster. A preset distance threshold ensures that clustering applied to different bundles yields a consistent density of streamlines. A minimum cluster size *C* is a hyperparameter such that any cluster smaller than *C* is pruned in this step. The bundle is then converted into a point-set representation for point cloud based Laplacian smoothing in the next step.

### 3.2 Step 2: Point cloud-based smoothing

While point clouds remove the sequential information in the streamlines, a smoothing operation applied to this representation can still consider a local neighborhood of points, which can be from neighboring streamlines. We assume that streamlines close to each other in a local neighborhood are more likely to share similar trajectories. One of the most common smoothing techniques, Laplacian regularization, has been successfully applied to point clouds [29],[30] using graph-based methods. In our work, the Laplacian matrix for a point cloud is computed using the approach described in Sharp and Crane (2022) [24]. They presented a robust replacement of the commonly used cotangent Laplacian by creating a ”tufted cover” for non-manifold edges and vertices and flipping edges to build Delaunay triangulations so that the Laplacian always has non-negative edge weights. This method can proves to be extremely robust, considering that point cloud triangulations contain many non-manifold edges or vertices. This approach is very fast, averaging 2.2 minutes to smooth point clouds ranging from 20K to 300K points in our analysis. Reconstruction of an example arcuate fasciculus with the Laplacian eigenfunctions is shown in Figure 2.

**Fig. 1.**
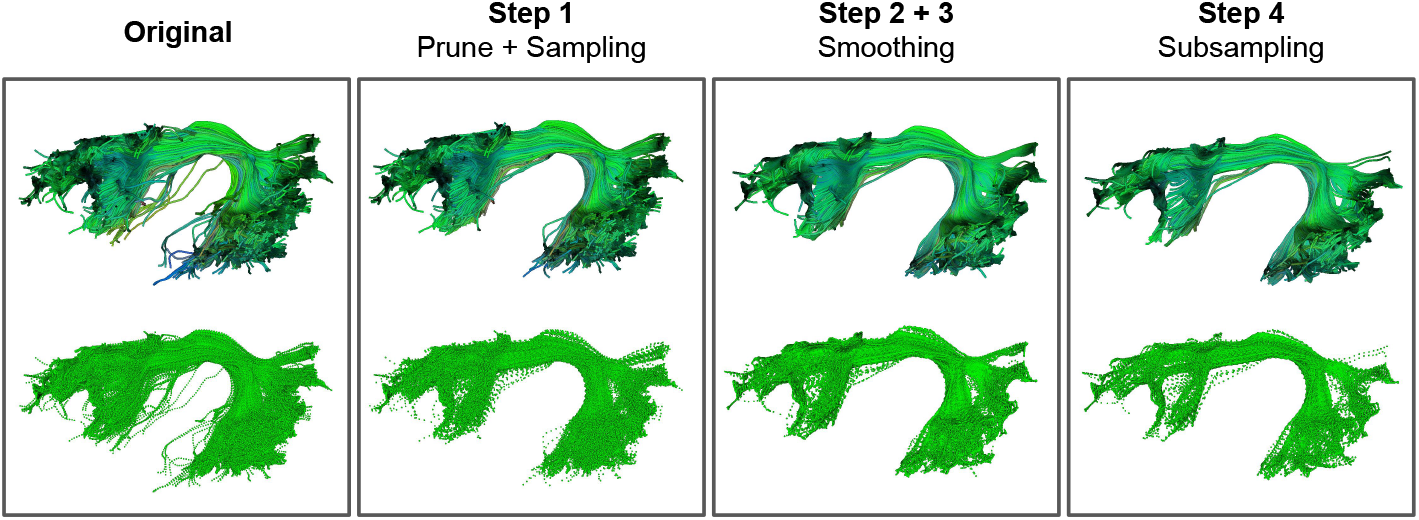
Here we show the 4 steps of *BundleCleaner* (see Section 3) on an example arcuate fasciculus bundle, comprised of two steps of pruning (steps 1 and 4), and two steps of smoothing (steps 2 and 3). Bundles are shown with both streamline (top) and point-cloud (bottom) representations.

**Fig. 2.**
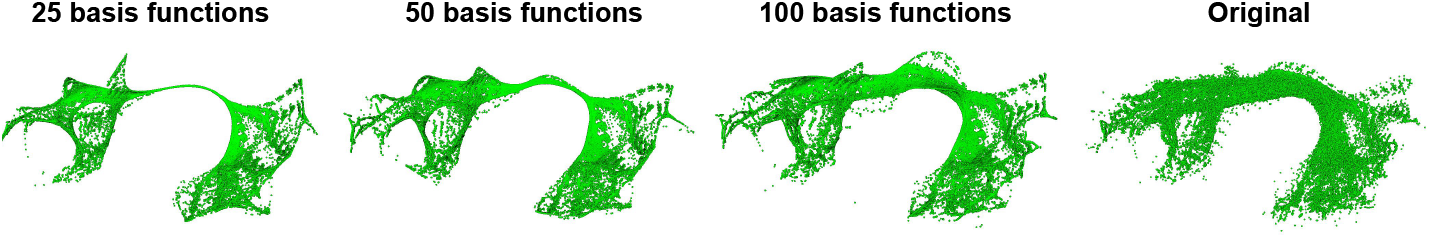
Point cloud based reconstruction of an arcuate fasciculus bundle with 25, 50, 100 eigenfunctions of the Laplacian matrix. The original bundle contains 62,253 points. The Laplacian matrix was built with a *K* = 100 sized neighborhood.

To compute the point cloud Laplacian, a local neighborhood of *K* neighbors is used to build a triangulation. The point cloud is then smoothed using Laplacian regularization,

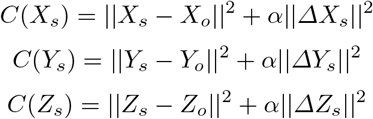

where *X*_*s*_,*Y*_*s*_, *Z*_*s*_ are the smoothed coordinates, *X*_*o*_,*Y*_*o*_, *Z*_*o*_ are the original co-ordinates, and *α* is the smoothing parameter, where larger *α* applies heavier smoothing. Each objective is smoothed using the conjugate gradient method.

### 3.3 Step 3: Streamline based smoothing

The bundle as a point cloud is reshaped back to streamlines for an additional step of smoothing using the Savitzky–Golay filter [23], an approach often used in 1D signal smoothing. The filter is applied to each streamline independently with second order polynomial and preset window size *W*, in case the denoised point clouds from Step 2 contain jagged edges. In addition, Sarwar *et al*. [22] have described the oscillating trajectories present in streamlines generated from probabilistic CSD as a *rippling* artifact, not present in those generated from deterministic methods. Higher *W* is therefore recommended for bundles generated from probabilistic tractography to produce smoother streamlines.

### 3.4 Step 4: Streamline-based subsampling

For efficient downstream tractometry analysis, BundleCleaner contains a step of streamline-based subsampling with a pruning procedure similar to Step 1 described in Section 3.1. While smoothing in Steps 2 and 3 would ideally remove and denoise spurious streamlines, extracted bundles can still contain streamlines that deviate locally in their orientation and curvature when the local neighborhood used in constructing the point-cloud Laplacian are all from outlier streamlines. The extra pruning in this step will further remove streamlines that could not be denoised. QuickBundles based clustering is applied to the output of Step 3 with the two parameters -distance threshold *D*^*′*^ and minimum cluster size *C*^*′*^ -similar to Step 1. The difference is that random subsampling is applied to each remaining cluster larger than the *C*^*′*^ with a subsampling rate of *S*.

## 4 Experiments & Results

We applied *BundleCleaner* to 8 bundles from each of the 66 participants -the left *arcuate fasciculus* (AF L), corpus callosum’s *forceps major* (CC ForcepsMajor), right *cingulum* (C R), left *corticospinal tract* (CST L), right *fronto-pontine tract* (FPT R), left *inferior longitudinal fasciculus* (ILF R), right *middle longitudinal fasciculus* (MdLF L), and left *uncinate fasciculus* (UF L) -with the following adjustments to the default parameters: *S*_*r*_ = 0.3, *α* = 150, *K* = 50, *W* = 10 and *S* = 0.75. *FiberNeat* [6] was applied to all 528 bundles using UMAP method^4^ for comparison with *BundleCleaner*. For bundles with fewer than 50 streamlines, only streamline smoothing (Step 3) is applied. Given the high levels of irregularities and the large number of streamlines generated from probabilistic fiber tracking [22], we resampled 30% points (*S*_*r*_ = 0.3) resulting in roughly 100 points per streamline on average, and apply more aggressive smoothing with a larger *α, K* and *W* value. After resampling, the distance between consecutive points in all streamlines remains smaller than the voxel size of 2mm, to ensure that dMRI-derived microstructural measures can be adequately mapped to points on the bundle. With *S*_*r*_ = 0.3, *S* = 0.75, and default pruning parameters, the resulting cleaned bundles contain 17% of points and 60% of streamlines compared to the original bundles on average. These downsampling factors were applied to allow data reduction to enable more efficient downstream analysis while retaining fidelity to the original tract geometry. To evaluate the results of *BundleCleaner*, we conducted a quantitative shape comparison and demonstrated how the cleaned bundle performs in localizing group differences in diffusion tensor imaging (DTI) measures using the BUAN tractometry pipeline.

### 4.1 Evaluation: Shape Comparison

We conduct shape comparison of the bundles before and after cleaning by computing two shape metrics -bundle *overlap* and *overreach* with respect to the atlas bundle [7]. We first voxelize each bundle *b* and the atlas bundle *a* using the same reference density map image into binary mask images *V*_*b*_ and *V*_*a*_, and then computing the fraction of non-zero voxels in *V*_*b*_ overlapping and overreaching with respect to *V*_*a*_.

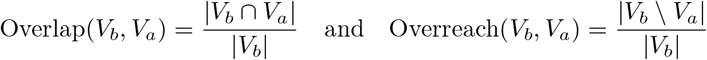

Figure 4 plots the two metrics for 8 bundle types before and after cleaning. We can see increased bundle overlap and decrease overreach scores in both cleaning methods, with *BundleCleaner* outperforming *FiberNeat* in particularly in C R, CST L and FPT R in both metrics. These results indicate that *BundleCleaner*, by incorporating smoothing can improve bundle alignment with the atlas bundles.

**Fig. 3.**
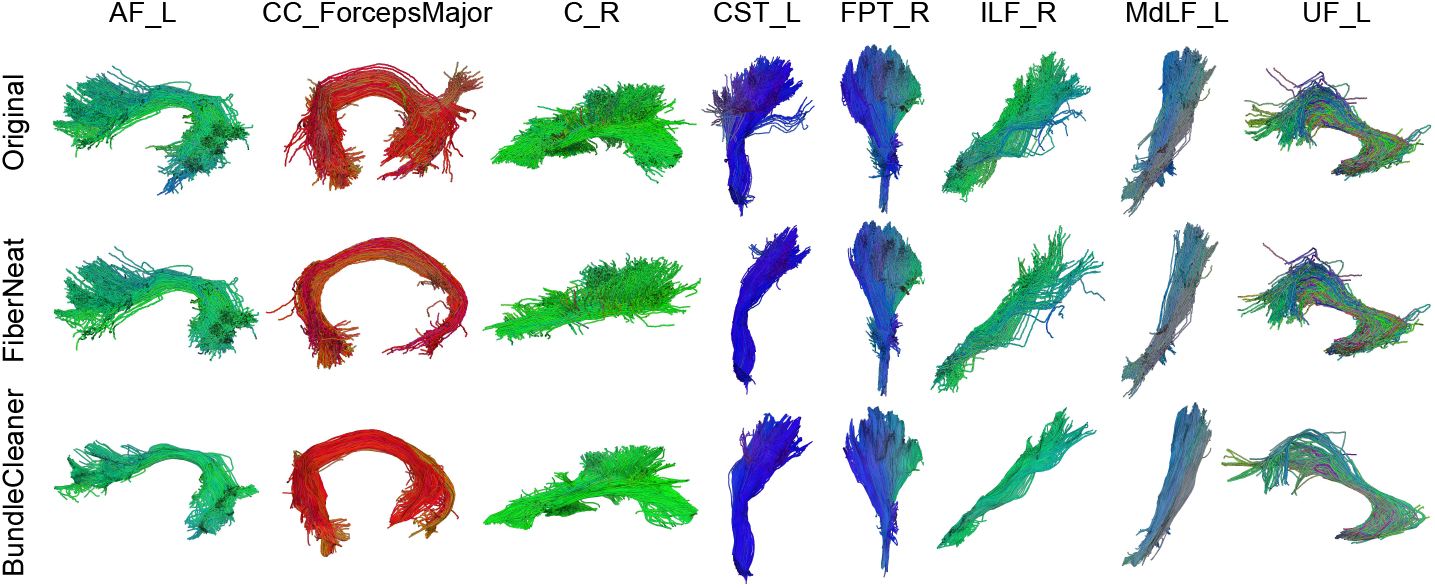
Examples of the 8 original, bundles cleaned with both *FiberNeat* and *Bundle-Cleaner*. Parameters for *BundleCleaner* are described in Section 4

**Fig. 4.**
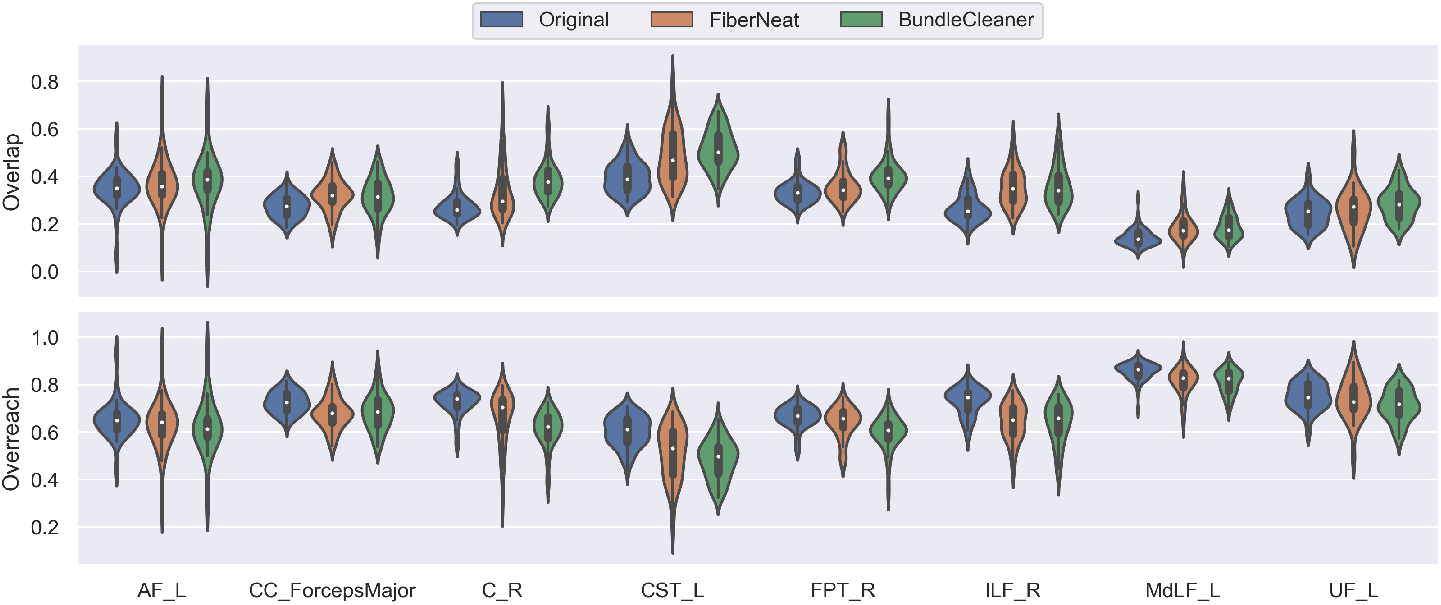
Shape comparison of the original, *FiberNeat* and *BundleCleaner* bundles, using metrics -bundle overlap and coverage with the atlas bundle defined in Section 4.1.

### 4.2 Evaluation: Tractometry

BUAN, using all points in a bundle, applies linear mixed models (LMM) to localize group differences along its length. We ran the BUAN tractometry pipeline on the aforementioned bundles from 32 CN versus 34 AD subjects, and investigated group differences in four DTI measures -fractional anisotropy (FA), mean diffusivity (MD), axial diffusivity (AD) and radial diffusivity (RD) -before and after cleaning. Fixed effect terms in the LMM include age and sex in addition to diagnostic group, and the random effect term is the subjects to account for multiple points belonging to the same subject in a segment being analyzed. False discovery rate (FDR) correction was applied to each bundle to account for multiple comparisons along the 100 tract segments.

Here, we show results for one of the DTI measures for each of six bundles in Figure 5 -plots on the left in each box show the negative logarithm of *p*-values on the *y*-axis, segments along the length of the bundle on the *x*-axis, along with FDR corrected threshold marked with a blue horizontal line. The standardized *β* values from the LMM are mapped onto the atlas bundle to indicate effect sizes on the right side of the box. The detected group differences are largely similar between the original and cleaned bundles -except for the end regions where shape variability is higher, and signals are less resistant to noise, notably in RD in segments 80-100 of the right cingulum. Other statistically significant differences after FDR correction are detected in the cleaned bundles -segment 10-30 in the right fronto-pontine tract, segment 20-40 in the left middle longitudinal fasciculus, and segment 30-40 in the left uncinate fasciculus -with similar if not larger effect size as reflected by the standardized *β*, despite using less than 20% of the points. This shows that *BundleCleaner* can robustly detect group differences in DTI measures while reducing the size of intermediate files by half.

**Fig. 5.**
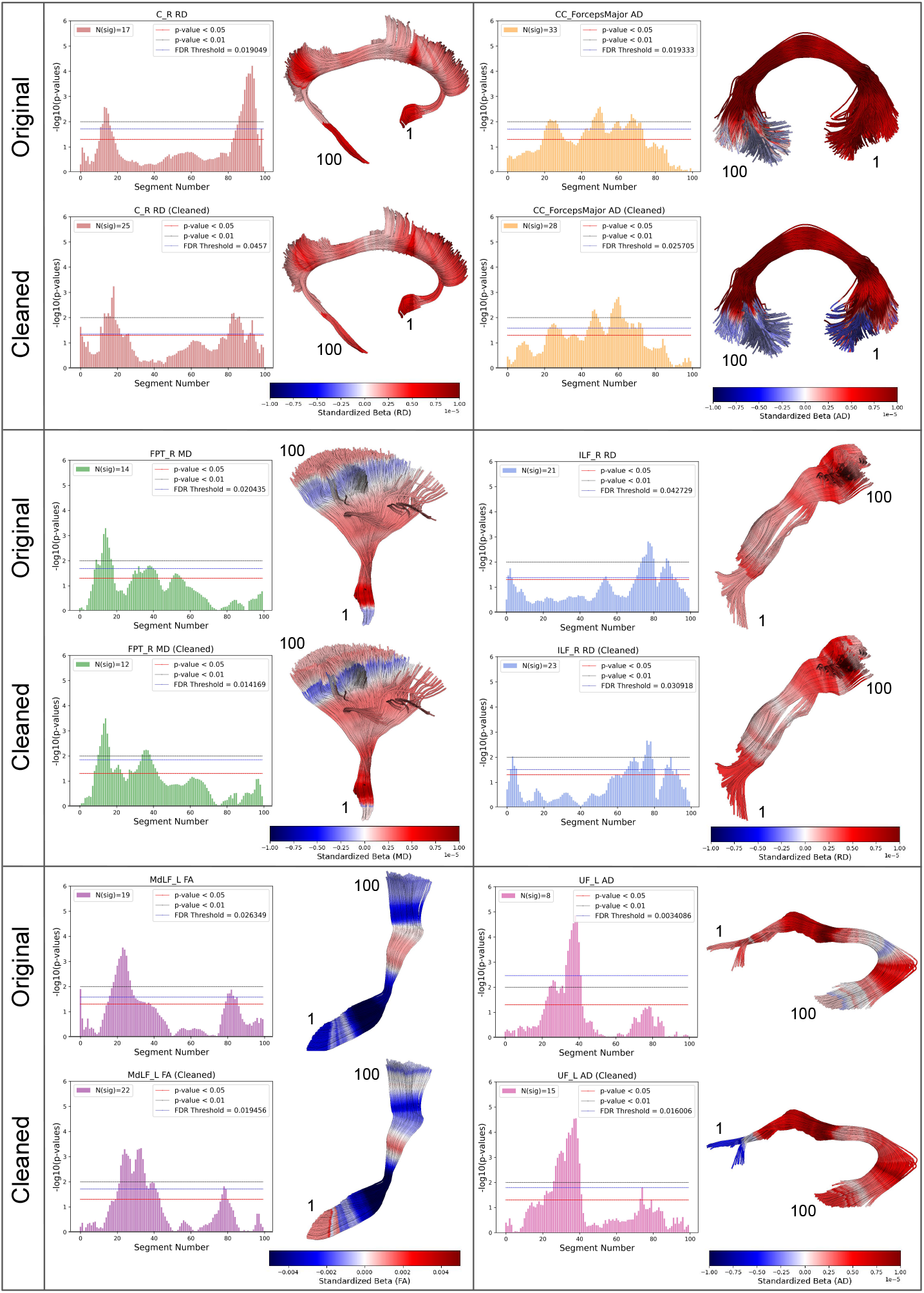
BUAN results for six bundles before and after cleaning. For each bundle, a bar plot of the *−* log(*p*) with the FDR threshold marked in blue (left), and standardized *β* mapped along 100 segments of the bundle (right) are shown. The group difference is shown between CN and AD participants. The number of segments with *p*-values that pass the FDR threshold is also marked in the bar plot.

## 5 Discussion

*BundleCleaner* parameters can be tuned to directly control global and local streamline smoothness as well as remove false positive. While a higher *α* value in Step 2 will produce ”tighter” bundles with more cohesion, a higher *W* value in Step 3 will produce streamlines with smoother trajectories; a higher minimum cluster size *C*/*C*^*′*^ in Step 1 and 4 will apply more aggressive streamline filtering. However, the approach relies on some degree of ‘correctness’ and overall cohesion of bundles produced by segmentation. In the case where a large proportion of the bundles is anatomically implausible, it is difficult for *BundleCleaner* to prune out such fibers. Figure 6 shows two examples in one CC ForcepsMajor, where cleaning was too aggressive, and one UF L bundle where it fails to remove a significant portion of false positive streamlines. We can reduce *C*/*C*^*′*^ for less aggressive pruning in the first case, and increase them or manually clean bundles using QuickBundles [8] for the second case. In addition, when cleaning bundles such as the corticospinal tract (CST) -where the lateral branch is likely to be removed - or bundles with large diameter but with sparse streamlines such as the corpus callosum, we recommend reducing *C*/*C*^*′*^ for a more lenient threshold.

**Fig. 6.**
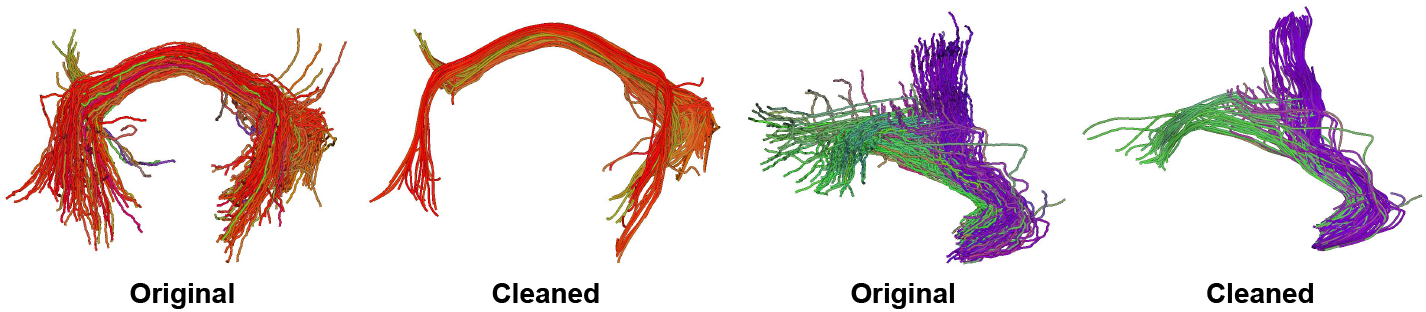
Two failure cases in one corpus callosum forceps major bundle and one left uncinate fasciculus bundle.

While each step of *BundleCleaner* can be run independently of the others, some parameters can be tuned together to provide better synergies. Resampling and pruning in step 1 ensure that fewer points are passed on to step 2, since point cloud-based smoothing is the most time-consuming step of this process. Lower *S*_*r*_ in Step 1 is therefore recommended for bundles generated with small tractography step sizes. Another effect of a lower resampling rate *S*_*r*_ is that it is more likely for points from neighboring streamlines to be considered in Laplacian smoothing, and this can be more directly tuned with *K* in Step 2. Lower *S*_*r*_ and higher *K* apply global bundle-based smoothing over a larger neighborhood while reducing the runtime in Step 2, and may be helpful for bigger and denser bundles.

*BundleCleaner* was developed with application to tractometry in mind and we take inspiration from smoothing operations in voxel-based morphometry prior to statistical analysis to increase the signal-to-noise (SNR) ratio [3]. While smoothing can remove finer fiber structure when applied to tractography data, we believe that the manually adjustable parameters in our method can offer value in practice for different use cases, such as tractometry and structural connectivity analysis. In our future work, we plan to test our approach on multi-shell datasets with higher b-values, streamlines generated with deterministic tractography, and bundles segmented from different atlases, to make corresponding parameter recommendations.

## 6 Conclusion

*BundleCleaner* is an unsupervised multi-step framework that can filter, denoise, and subsample bundles using both point cloud and streamline-based methods, which considers the global bundle structure as well as local streamline-wise features. This approach is efficient, can denoise bundles with more than 1 million points, reduce bundle overreach and improve alignment with the atlas bundles used in segmentation, making it easier to work with unwieldy tractography data in downstream analysis. We show in a downstream tractometry analysis that cleaned bundles represented with less than 20% of the points compared to the original bundles can robustly localize microstructural differences between CN and AD subjects in an Indian cohort, while significantly reducing memory burden. This approach shows promise for large-scale multi-site tractometry analysis. *BundleCleaner* parameters are intuitive to tune and can be applied to any fiber bundles with varying noise levels.

## Footnotes

4 FiberNeat implementation is available at https://github.com/BramshQamar/FiberNeat/tree/main

